# Aberrant Cortical Activity In Multiple GCaMP6-Expressing Transgenic Mouse Lines

**DOI:** 10.1101/138511

**Authors:** Nicholas A. Steinmetz, Christina Buetfering, Jerome Lecoq, Christian R. Lee, Andrew J. Peters, Elina A. K. Jacobs, Philip Coen, Douglas R. Ollerenshaw, Matthew T. Valley, Saskia E. J. de Vries, Marina Garrett, Jun Zhuang, Peter A. Groblewski, Sahar Manavi, Jesse Miles, Casey White, Eric Lee, Fiona Griffin, Joshua D Larkin, Kate Roll, Sissy Cross, Thuyanh V. Nguyen, Rachael Larsen, Julie Pendergraft, Tanya Daigle, Bosiljka Tasic, Carol L. Thompson, Jack Waters, Shawn Olsen, David J. Margolis, Hongkui Zeng, Michael Hausser, Matteo Carandini, Kenneth D. Harris

## Abstract

Transgenic mouse lines are invaluable tools for neuroscience but as with any technique, care must be taken to ensure that the tool itself does not unduly affect the system under study. Here we report aberrant electrical activity, similar to interictal spikes, and accompanying fluorescence events in some genotypes of transgenic mice expressing GCaMP6 genetically-encoded calcium sensors. These epileptiform events have been observed particularly, but not exclusively, in mice with Emx1-Cre and Ai93 transgenes, across multiple laboratories. The events occur at >0.1 Hz, are very large in amplitude (>1.0 mV local field potentials, >10% df/f widefield imaging signals), and typically cover large regions of cortex. Many properties of neuronal responses and behavior seem normal despite these events, though rare subjects exhibit overt generalized seizures. The underlying mechanisms of this phenomenon remain unclear, but we speculate about possible causes on the basis of diverse observations. We encourage researchers to be aware of these activity patterns while interpreting neuronal recordings from affected mouse lines and when considering which lines to study.

## Introduction

A paramount goal for neuroscience is the development of technologies that can measure neural activity across large regions of the brain with high temporal and spatial resolution. The development of calcium-sensitive fluorescent proteins has allowed for the minimally-invasive measurement of neuronal activity with high signal-to-noise, at single neuron resolution, and on increasingly large spatial scales (Chen et al., 2013; Kim et al., 2013; Sofroniew et al., 2016; Stirman et al., 2016). Genetically encoded calcium-sensitive fluorescent proteins offer a number of advantages over viral expression or bulk injection: the experimental procedures can be less invasive; expression levels are more consistent among cells and across the subject’s lifetime; and they allow for imaging across wider regions of the brain. Accordingly, they have become popular tools and several versions of this technology have been developed, in which different versions of the sensor protein are expressed under various promoters (Dana et al., 2014; Madisen et al., 2015; Wekselblatt et al., 2016).

Here we compare the neural activity, measured electrophysiologically and with imaging, from a variety of transgenic mouse lines and find that a subset of these lines exhibit aberrant activity patterns resembling interictal spikes. Due to this resemblance, discussed further below, we refer to these events as “epileptiform”. We describe the characteristics of this activity, its incidence across different mouse lines, and speculate about its possible causes.

## Results

### “Epileptiform” events in local field potential (LFP) recordings

We observed highly distinctive, aberrant electrophysiological events in LFP recordings from isocortex of some GCaMP-expressing lines. Epileptiform events were seen in all recorded Emx1-Cre;Camk2a-tTA;Ai93 mice, but not in mice from several other lines, including wild-type C57BL/6J (Figure 1). Events were characterized by large amplitude, brief duration, stereotyped shape within a recording, and characteristic periodic spacing (“refractory period” of ~0.5 - 1sec, overall rate ~0.1 – 0.5Hz). The presence of the events was visualized using a scatter plot of the prominence and width of all negative LFP peaks (see Methods). In recordings exhibiting these events, this plot revealed a distinct cluster of events of high prominence and short duration, outside the range of variation observed in other genotypes. Though the shapes of the events were stereotyped within a recording, they could vary significantly across recordings, presumably reflecting differences in the location of the recording site and reference relative to the origin and spatial spread of the event at a larger spatial scale (see also imaging results, below). These events were larger in amplitude than individual action potentials (>1mV versus <750µV), longer in duration (>10ms versus <2ms), and visible over a greater spatial range (>1mm versus <100µm), so they cannot reflect the activity of single neurons.

**Figure 1:**
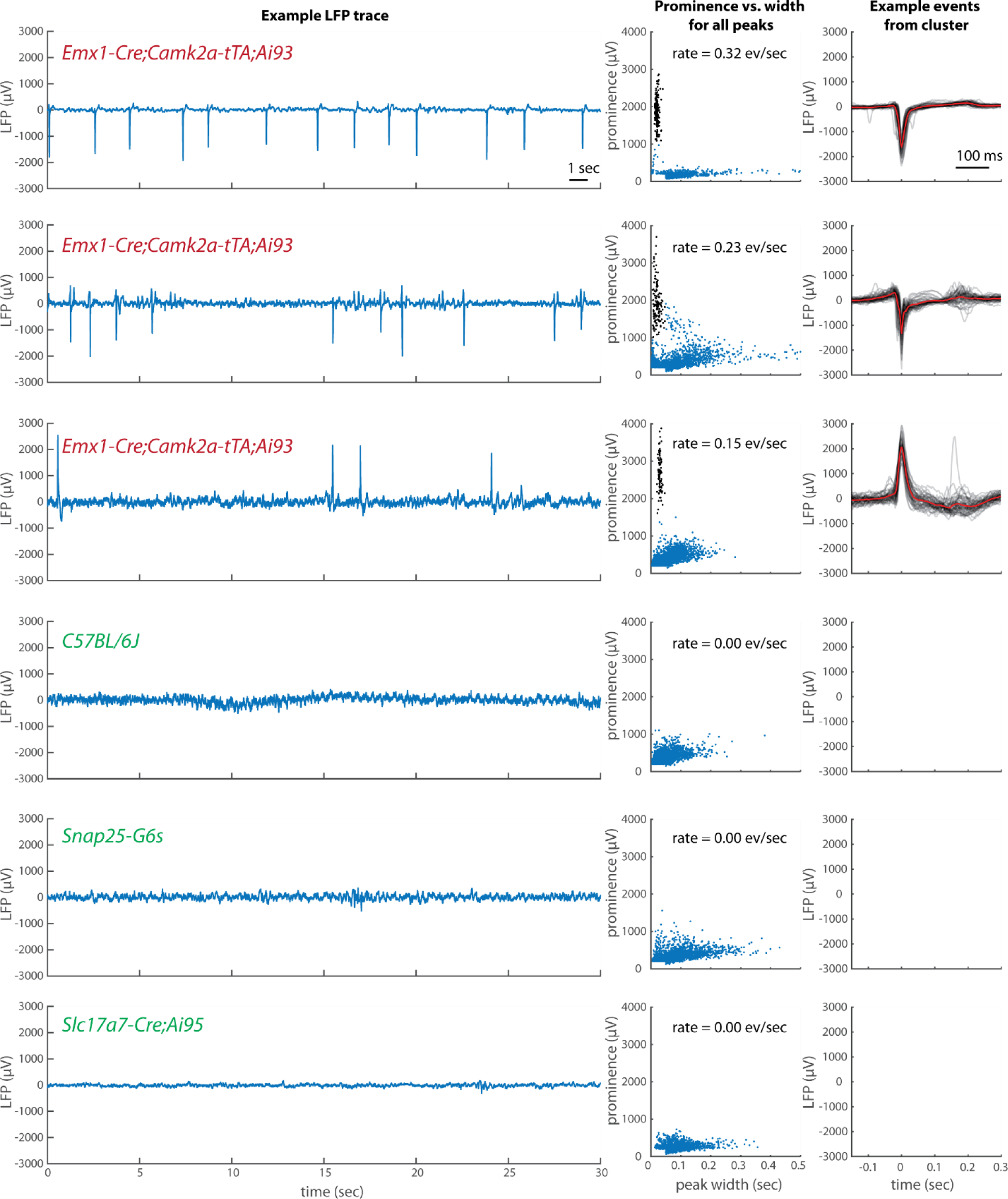
Epileptiform electrical activity in frontal cortex of some GCaMP6-expressing mice but not others. Left, each row contains an example segment of raw LFP data from each of six mice, with genotype identified in colored text. Middle, a plot of the prominence vs. width of all peaks (see Methods) in the full LFP traces. Points highlighted in black were identified as a distinct cluster and included in the computation of event rate and the example events plotted at right. Right, 50 example events (grey) and the mean of all events (red). For the recording in row C, positive peaks rather than negative were analyzed.

We tested for epileptiform events in electrophysiological recordings from ten transgenic mice with six different combinations of transgenes, plus one wild-type C57BL/6J mouse (Table 1). All tested Emx1-Cre;Camk2a-tTA;Ai93 mice (4/4) showed these events. However, we did not observe these events in mice of four other transgenic lines (Slc17a7-Cre;Ai95, Camk2a-tTA;tetO-G6s, Snap25-G6s, and PV-Cre;Ai32) or in the wild-type mouse. A single recorded Emx1-Cre;Camk2a-tTA;Ai94 mouse showed these events, but as this mouse was selected for this recording based on the presence of epileptiform events in imaging, their presence in electrophysiology in this mouse does not indicate that all mice of that genotype exhibit them.

**Table 1:** Incidence of epileptiform events in electrophysiological recordings of local field potentials in cortex. Each count represents one mouse recorded electrophysiologically. *The Emx1-Cre;Camk2a-tTA;Ai94 mouse was selected for electrophysiology on the basis of epileptiform events observed in imaging.

| Genotype | Incidence of events |
| --- | --- |
| Emx1-Cre;Camk2a-tTA;Ai93 | 4/4 mice |
| Emx1-Cre;Camk2a-tTA;Ai94 | 1/1* |
| Slc17a7-Cre;Ai95 | 0/1 |
| Camk2a-tTA;tetO-G6s | 0/1 |
| Snap25-G6s | 0/1 |
| Pvalb-Cre;Ai32 | 0/2 |
| wildtype (C57BL/6J) | 0/1 |

### Epileptiform events in widefield imaging

Epileptiform events could also be unambiguously identified with widefield imaging, using the same characteristics, i.e. large amplitude and brief duration relative to other events. In an example mouse the rate of occurrence, refractory period, relative amplitude, and relative incidence across cortical regions were consistent, suggesting that the >1 mV events observed in LFP were the same events identified in imaging (Figure 2). This was also true in other mice with non-simultaneous measurements of LFP and imaging. We conclude that the large, brief “flashes” observed in imaging correspond to the large, brief LFP events.

**Figure 2:**
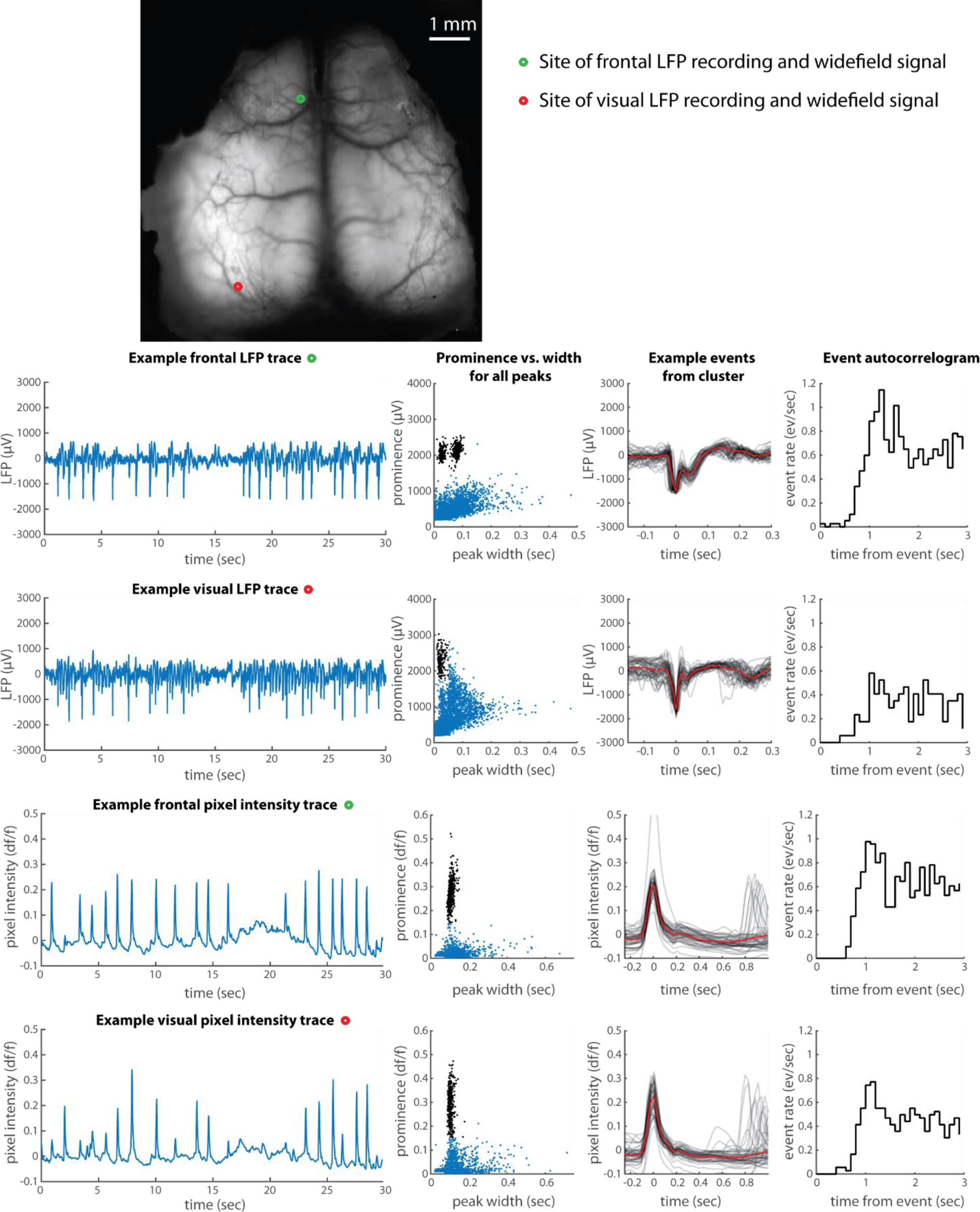
Comparison of events observed in LFP and widefield calcium imaging in the same mouse. Figure format as in Figure 1. The two LFP recordings were made simultaneously with each other, but not simultaneously with the widefield imaging. In the frontal LFP recording, the two clusters in prominence vs. width arise because of the double-peaked shape of the events in this mouse at this site: for some events the width includes only the first peak, for some it includes both.

By observing the epileptiform events in widefield imaging, we were able to measure the spatial extent of the events and to characterize their incidence over a wider range of mice, including multiple genotypes and mice from multiple institutions. Widefield data from six more example mice are shown in Figure 3. The events were most frequently observed and relatively largest in an antero-lateral region of cortex including somatosensory, motor, and frontal cortex. Nevertheless they could on some occasions be observed in visual cortex or retrosplenial cortex, usually with lower amplitude and/or relatively infrequent occurrence. They were nearly always bilaterally symmetric.

**Figure 3:**
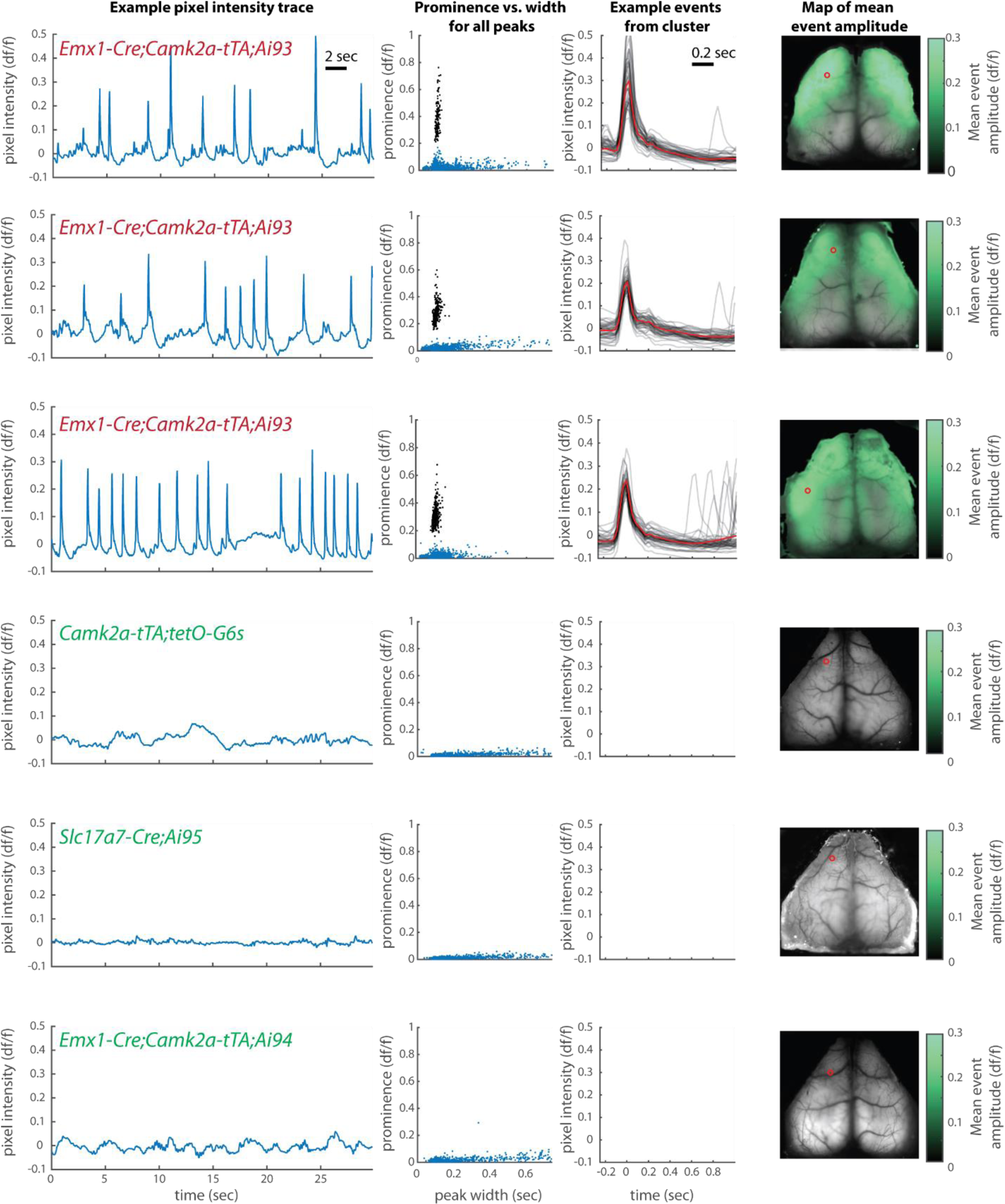
Incidence and spatial extent of epileptiform events observed in widefield calcium imaging. Plot format as in Figure 1. The intensity trace and detected peaks come from the pixel indicated by the red circle on the brain image at right. Green coloration in the right-most panel, overlaid on the mean image of the brain, represents the amplitude of the mean event at each point across the brain.

### Incidence

We observed epileptiform events in widefield imaging data from 61 of 138 mice across 15 genotypes and four labs (Table 2). These observations were made either using limited windows over restricted regions of cortex or, in separate mice, full-hemisphere windows. The incidence of events was greater in full-hemisphere recordings for matched genotypes. Considered together with the observations above that the epileptiform events often did not travel to visual cortex or had substantially lower amplitude there, we infer that they were often undetectable when limited regions of cortex were imaged alone.

**Table 2:** Incidence of epileptiform events as observed with widefield calcium imaging. Epileptiform events were judged to occur by manually inspecting raw imaging movies, traces, and prominence versus width plots for peaks of the traces, as shown in Figure 3. See Methods for details of mouse lines and imaging preparations. Red, yellow, and green coloration indicates the relative incidence of events (high, rare, or never observed). All mice in this table either had intact Cre conditionality or it was unknown. Camk2a* indicates that these mice were treated with doxycycline until 7 weeks (see below).

| Institute/Lab | Cre genotype | tTA genotype | GCaMP genotype | Incidence of events in any size imaging window | Incidence of events in full-hemisphere imaging |
| --- | --- | --- | --- | --- | --- |
| UCL-Carandini/Harris | Emx1 | Camk2a | Ai93 | 12/12 mice | 12/12 mice |
| Allen Institute |  |  |  | 8/19 | 5/7 |
| Rutgers-Margolis | Emx1 | Camk2a | Ai93 | 3/3 | 3/3 |
| Rutgers-Margolis | Emx1 | Rosa26 | Ai93 | 7/7 | 7/7 |
| UCL-Hausser | Emx1-Kess | Camk2a | Ai93 | 7/12 | - |
| Allen Institute | Slc17a7 | Camk2a | Ai93 | 1/5 | 1/5 |
| Allen Institute | Cux2 | Camk2a | Ai93 | 1/5 | - |
| UCL-Carandini/Harris | Emx1 | Camk2a | Ai94 | 1/5 | 0/4 |
| Allen Institute | Rbp4 | Camk2a | Ai93 | 0/11 | - |
| Allen Institute | Rorb | Camk2a | Ai93 | 0/3 | - |
| Allen Institute | Scnn1a | Camk2a | Ai93 | 0/8 | - |
| UCL-Hausser | Emx1-Kess | Camk2a* | Ai93 | 0/15 | - |
| UCL-Carandini/Harris | Slc17a7 | - | Ai95 | 0/4 | 0/4 |
| Allen Institute | Emx1 | - | Ai95 | 0/1 | 0/1 |
| Allen Institute | Emx1 | - | Ai96 | 0/3 | 0/3 |
| UCL-Carandini/Harris | - | Camk2a | tetO-G6s | 0/6 | 0/6 |
| Allen Institute |  |  |  | 0/6 | 0/2 |
| UCL-Carandini/Harris | - | - | Snap25-G6s | 0/4 | 0/4 |
| Allen Institute |  |  |  | 0/2 | 0/2 |
|  |  |  |  | High incidence |  |
|  |  |  |  | Low incidence |  |
|  |  |  |  | No events observed |  |

Epileptiform events were seen in the great majority of mice with the Emx1-Cre, Ai93, and either Camk2a-tTA or Rosa26-tTA transgenes (31/33 mice with full-hemisphere recordings; 44/59 in total). They were also seen in mice that expressed Ai93 with other drivers, and in which germline Cre recombination had occurred (15/22 mice with limited-size windows, versus 0/14 in the same lines when germline Cre recombination did not occur; Table 3). Germline Cre recombination is the phenomenon where ectopic expression of Cre in germline cells of previous generations results in deletion of the Lox-stop-Lox upstream of the reporter, and thus results in a loss of Cre conditionality in all cells of the animal (Schmidt-Supprian and Rajewsky, 2007). In these cases, the GCaMP expression pattern is determined by tTA expression alone, regardless of the cell types expressing Cre (Figure 4; in all such cases, tTA expression was driven by the Camk2a promoter, resulting in broad and dense expression). In our experiments, we observed germline Cre recombination specifically with Emx1-Cre, Ntsr1-Cre, Rbp4-Cre, and Rorb-Cre. Although not all mice were tested for germline recombination, we observed a high probability of epileptiform events in Emx1-Cre;Camk2a-tTA;Ai93 mice with Cre conditionality intact (i.e., germline Cre recombination had been confirmed absent with genotype tests; 17/19 full-hemisphere recordings, 20/31 in total). We therefore conclude that mice with germline Cre recombination have a high probability of having epileptiform events for all tested Cre driver lines, but that the Emx1-Cre line is a special case in which epileptiform events were common regardless of germline recombination.

**Table 3:** Effects of germline Cre recombination on incidence of epileptiform events. The column of observations “WITH germline Cre recombination” indicates mice that had lost Cre conditionality, whereas the column “WITHOUT” indicates mice that were normal, i.e. expressed GCaMP only in subsets of neurons according to the Cre driver line employed. In this table, all mice were imaged with widefield calcium imaging of any size window.

| Cre genotype | tTA genotype | GCaMP genotype | Incidence WITH germline Cre recombination | Incidence WITHOUT germline Cre recombination |
| --- | --- | --- | --- | --- |
| Emx1 | Camk2a | Ai93 | 9/10 | 20/31 |
| Ntsr1 | Camk2a | Ai93 | 6/10 | - |
| Rbp4 | Camk2a | Ai93 | 4/4 | 0/11 |
| Rorb | Camk2a | Ai93 | 5/8 | 0/3 |

**Figure 4:**
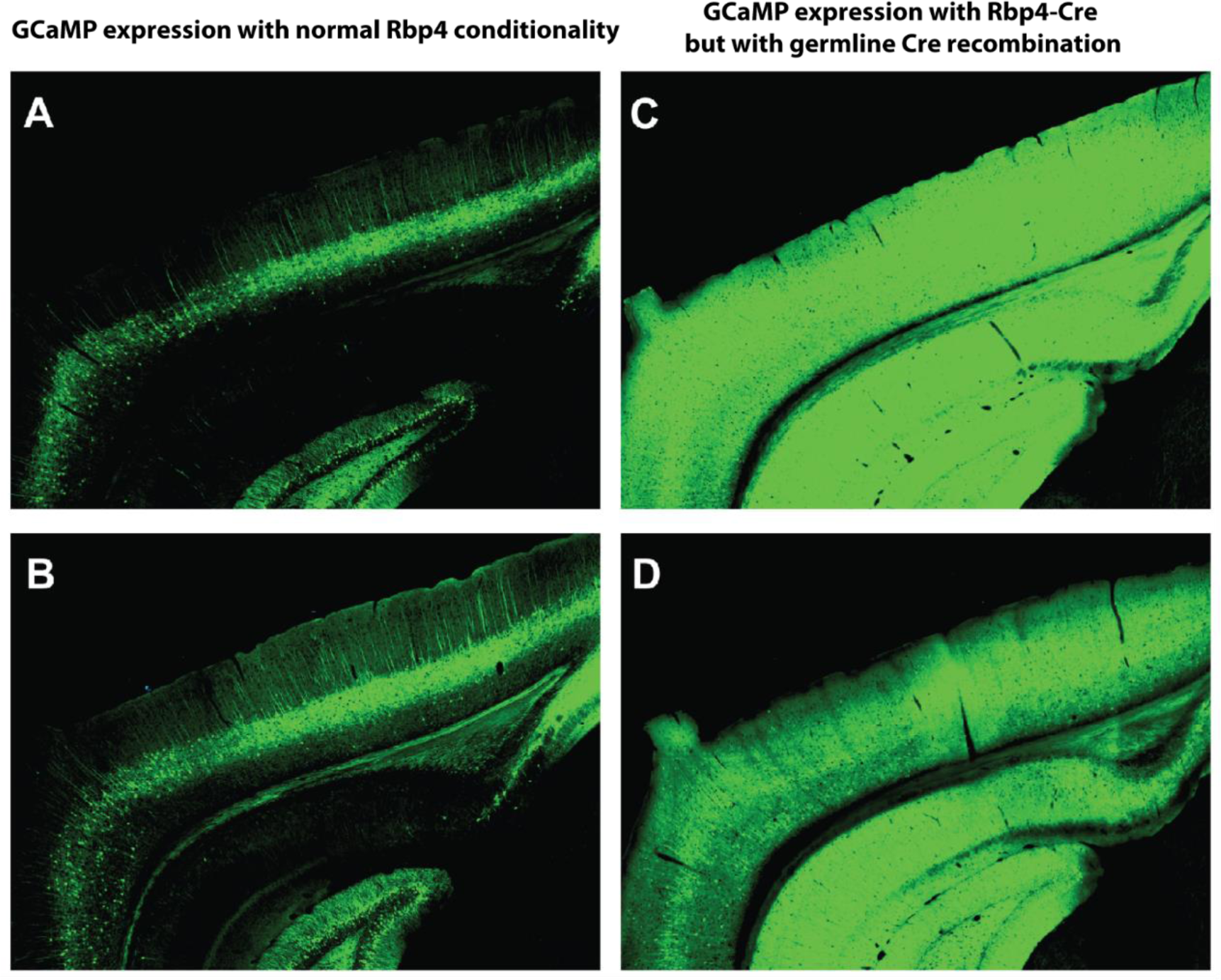
Germline Cre recombination results in widespread GCaMP expression. A, B, Native GCaMP6 fluorescence obtained using 2-photon serial tomography from two Rbp4-Cre/wt;Camk2a-tTA/wt;Ai93 Stop+/Ai93 Stop+ mice, showing expression of GCaMP only in restricted populations of L5 cortical neurons and moderate expression in hippocampus. C, D, Similar images with matched intensity scale but from a Rbp4-Cre/wt;Camk2a-tTA/wt;Ai93 Stop+ / Ai93 Stop deleted mouse that had germline Cre recombination, showing high, widespread expression across cortex and hippocampus.

Only three mice with confirmed absence of germline recombination and genotype not including both Emx1-Cre and Ai93 were observed to have epileptiform events. Their genotypes were Cux2-Cre;Camk2a-tTA;Ai93 (1/5 mice recorded with limited-size windows), Emx1-Cre;Camk2a-tTA;Ai94 (1/5 mice overall, 0/4 with full-hemisphere imaging), and Slc17a7-Cre;Camk2a-tTA;Ai93 (1/5 mice recorded with full hemisphere windows). The other eight genotypes tested had zero observations of these events (0/48 mice).

### Detection of epileptiform events in 2-photon imaging

The epileptiform events were also detectable in 2-photon calcium imaging by creating a trace of the mean activity across the entire imaging field of view, including neurons and neuropil. In one mouse we measured the neural activity non-simultaneously in all three modalities (LFP, widefield, 2-photon) and found the same signatures in all cases (Figure 5). We conclude that epileptiform events can be detected in at least some cases when present in the 2-photon field of view. The caveat described above - that epileptiform events can be missed when limited fields of view are employed that do not include the extent of the events - applies especially to 2-photon imaging experiments with small fields of view, and for this reason we have not summarized in detail our observations with 2-photon imaging.

**Figure 5:**
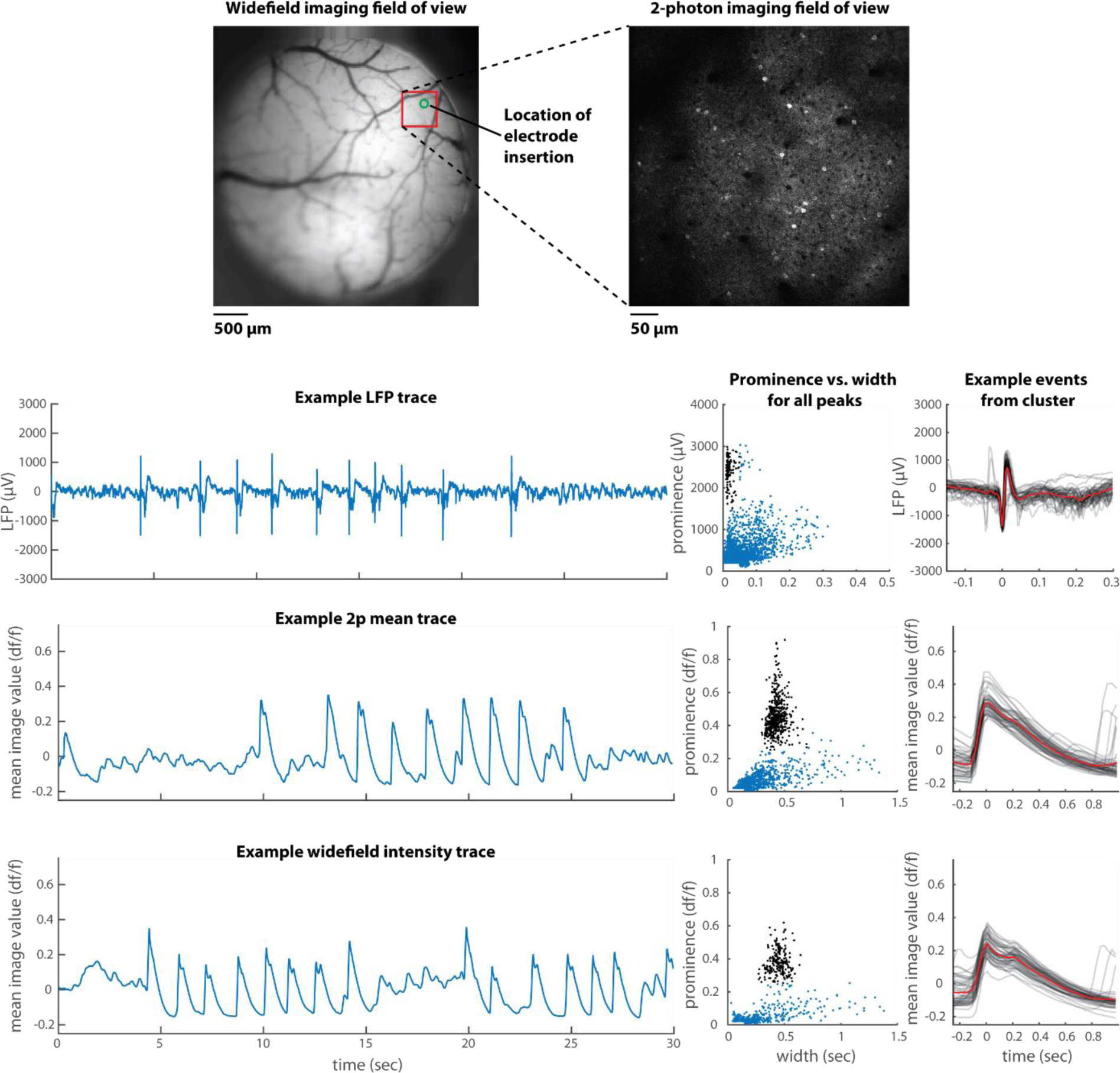
Epileptiform events observed in LFP, 2-photon calcium imaging, and widefield calcium imaging in one individual mouse, but not simultaneously. Plot formats as in Figure 1. The genotype of the mouse was Emx1-Cre;Camk2a-tTA;Ai94 (expressing GCaMP6s). 2-photon trace was generated as the mean intensity of each frame across the entire field of view; widefield trace was generated as the mean within an ROI approximating the 2-photon field of view.

### Generalized seizures

We observed generalized, tonic-clonic type seizures in 11 mice at three laboratories, including twice during a widefield imaging session. Of these, 10 were Emx1-Cre;Camk2a-tTA;Ai93 genotype or had germline Cre recombination with a different Cre line. These 10 mice comprise a small proportion of the total number of such mice that were studied (10/103, 9.7%), but this seizure incidence still represents a significant increase relative to other genotypes (1/68, 1.5%; p<0.01, binomial test; the one mouse with a reported seizure in this group was Scnn1a-Cre;Camk2a-tTA;Ai93). However, mice were not monitored constantly in their home cages so these numbers reflect a lower bound on the proportion of mice that had seizures.

### Prevention of events with doxycycline treatment

To test the role of GCaMP6 expression during development, we treated a cohort of Emx1-Cre-Kess;Camk2a-tTA;Ai93 (n=15) and Emx1-Cre-Kess;Camk2a-tTA;Ai94 (n=2) mice with doxycycline from birth until aged 7 weeks. Doxycycline blocks tTA activity and therefore prevented expression of GCaMP6 until removal from the drinking water. GCaMP6 expression levels were low shortly after removal of doxycycline and ramped up to levels comparable to untreated Emx1-Cre;Camk2a-tTA;Ai93 within 3-4 weeks. Using widefield imaging above S1, we found that all mice treated with doxycycline were free of epileptiform events in this brain area up until at least 20 weeks of age, though one mouse did develop seizures despite lack of detected events. To compare this to the onset of epileptiform activity in untreated mice we performed longitudinal widefield imaging in mice starting at the age of 7 weeks (n=2), 8 weeks (n=1) or 11 weeks (n=2). In these mice epileptiform events were not present at 8 weeks of age (n=0/3) but developed by the age of 13 weeks (n=5/5 imaged at 13 weeks or greater; Figure 6). We conclude that doxycycline treatment delays or reduces the occurrence of epileptiform events.

**Figure 6:**
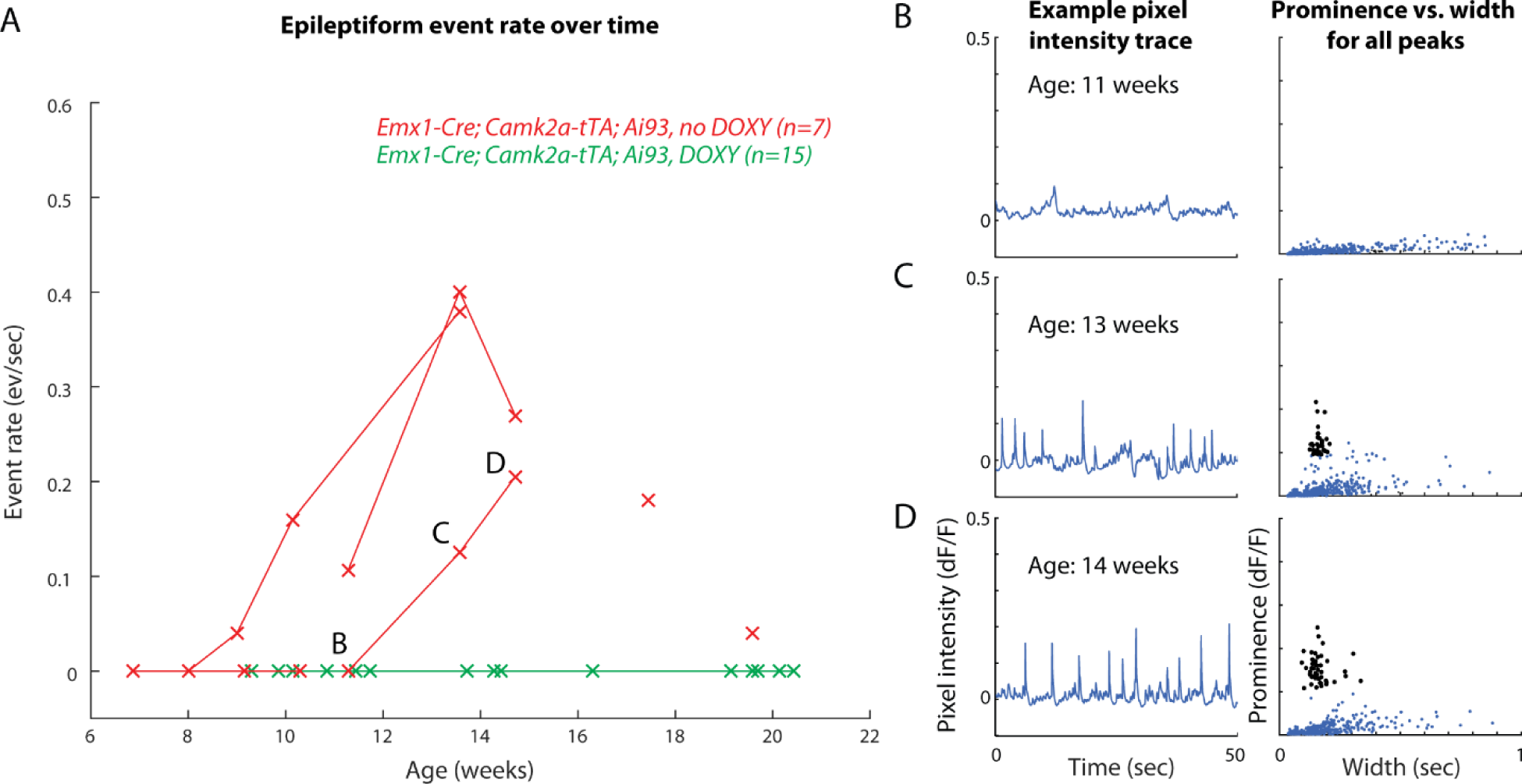
Epileptiform events fail to develop over time in mice treated with doxycycline until age 7 weeks. A, Measured rates of epileptiform events in control (red) and doxycycline treated (green) mice by age at time of measurement. Connected points indicate observations from the same mouse. All doxycycline-treated mice failed to develop events over the measured time period. Note that the mice represented in this figure are a small subset of all measured mice and we do not intend to claim that the time courses followed here are representative of all mice developing events. Letters B, C, and D refer to the measurements corresponding to the figure panels at right. B-D, example traces and prominence vs. width plots for an example control mouse showing development of events between 11 and 14 weeks.

## Discussion

Here we report the observation of aberrant neural activity resembling interictal spikes in some genotypes of transgenic mice expressing GCaMP6. This activity was most prominent in the Emx1-Cre;Camk2-tTA;Ai93 genotype, or in Camk2a-tTA;Ai93 mice that had undergone germline Cre recombination, but was also occasionally observed in other genotypes including one where GCaMP expression was driven by the Ai94 rather than Ai93 line.

These epileptiform events are strikingly distinct from activity observed in unaffected mice, in both amplitude and time course. Nevertheless, they can be easy to miss, for several reasons. First, there are no obvious behavioral manifestations, other than very rare generalized seizures. Second, the events are typically absent in visual cortex, a commonly studied area. Finally, the events may be obscured in 2-photon imaging, even when present in the field of view, either because the sampling rate is slow relative to their duration or because they may primarily appear in neuropil and many individual neurons may show little relationship to them (e.g., as shown in Muldoon et al., 2015).

The presence of this aberrant activity complicates interpretation of studies on such mice. Certainly, when aberrant events are present in the recorded neuronal tissue, care must be taken when generalizing any conclusions on brain function drawn from these recordings to other mouse genotypes. For example, as the epileptiform events were substantially larger than other observed activity, they dominate measurements such as response variability and correlations across regions. Furthermore, our results suggest that the epileptiform events in these mice may frequently be undetectable in visual cortex even though present in other brain regions. In such cases, it is possible that visual cortical activity shows more subtle alterations, not revealed by the present analyses.

Despite the large amplitude of these events, performance in learning and executing trained behavioral tasks was not immediately distinguishable between mice with and without them, and visual responses were similar to those in wild-type mice, when periods containing aberrant events were excluded (data not shown). Tonic-clonic seizures were also rare in these mice. Therefore, it is possible that some properties of cortical functional organization may still be measured with imaging in these mice, such as receptive field maps (Zhuang et al., 2017).

### Relationship to interictal spikes

In epileptic patients and animal models, interictal spikes are a commonly observed phenomenon. These large, brief electrical events are observable in EEG, ECoG, and intracranial recordings, and are considered to always be pathological (Rodin et al., 2009). Whether they merely represent a symptom of a damaged brain, actively induce future seizures, or even act preventatively remains disputed (Staley et al., 2011). Their exact characteristics vary by species and by variety of epilepsy, but in mice they have been defined as having <200ms duration and large amplitude (generally >2x background activity amplitude, e.g., Erbayat-Altay et al., 2007). In these ways they are similar to the aberrant events we have observed, and we therefore referred to them as “epileptiform”. Furthermore, we observed a small proportion of mice to have generalized seizures among the mice with genotypes linked to a high incidence of epileptiform events. For these reasons, it seems possible or likely that the events we observed are similar or identical in origin to interictal spikes, and that the mice with these events may indeed be epileptic. To establish this more firmly, one would need 24-hour observation of mice to establish seizure incidence, as well as a clearer understanding of the mechanisms underlying the events we observed. Note that while the epileptiform events observed here appear similar to interictal spikes, they are dissimilar to the spike-wave discharges noted in many rat strains and some mouse models of epilepsy, which occur at a much higher rate of 7-9Hz (Wiest and Nicolelis, 2003; Fisher et al., 2014).

### Cause of aberrant activity

The cause of the epileptiform events is yet unknown and may result from a combination of effects, including Cre toxicity, tTA toxicity, and genetic background. Considering all observations together, however, the broad expression of GCaMP itself, particularly during development, seems likely to be a major factor.

The Cre enzyme is toxic (Schmidt-Supprian and Rajewsky, 2007) and Emx1-Cre specifically causes enhanced seizure susceptibility (Kim et al., 2013). Indeed we did not observe epileptiform events in any of the tested mice lacking Cre expression (i.e. Snap25-G6s, Camk2a-tTA;tetO-G6s, and wild type C57BL/6J, n=0/19 mice), consistent with a potential contributing role of Cre. However, regarding Emx1-Cre, we observed the events also in mice with Slc17a7-Cre, Ntsr1-Cre, Rbp4-Cre, and Rorb-Cre, so this particular Cre expression pattern cannot alone explain the effects.

tTA can also be neurotoxic, causing hippocampal degeneration in mice of at least some strains, including 129X1/SvJ (related to the strain of origin for Ai93 and Ai94, 129S6/SvEvTac, which was not tested; Han et al., 2012). In addition to the interaction with tTA toxicity, strain itself might play a role as different strains of mice have different seizure susceptibility, and the 129S3/SvImJ strain (again related but not identical to the strain of origin for Ai93 and Ai94) has higher seizure susceptibility than some other strains including C57BL/6J (Frankel et al., 2001). In general, the mice in this study were not congenic to a C57BL/6J background. However, the Rbp4-Cre mice with versus without germline Cre recombination have identical genetic backgrounds and identical levels of Cre and tTA, yet show large differences in event incidence (also true for Rorb-Cre). This comparison rules out a determining contribution of tTA toxicity or genetic background, though the possibility of an interaction remains.

GCaMP expression itself may play a role in the genesis of the epileptiform events. GCaMP binds calcium and thus buffers its intracellular concentration. Calcium plays many important roles in neurons, for example in synaptic transmission and in the expression of genes for synaptic plasticity, and disrupting these roles may accordingly alter network activity. Consistent with this, some genetic models of epilepsy in mice result from mutations to calcium channels (Letts et al., 1998). A major (if not the only) difference between the Rbp4-Cre;Camk2a-tTA;Ai93 mice with versus without germline Cre recombination is that the former will express GCaMP in a larger subset of neurons and possibly at higher levels than the latter, and the major difference in event incidence between these two groups of subjects (also true for Rorb-Cre) strongly suggests that the GCaMP expression itself plays a role.

Finally, we tested the role of GCaMP expression during development by suppressing tTA activity with doxycycline until 7 weeks postnatal in a cohort of mice. This treatment was effective in eliminating epileptiform events while preserving high expression levels in adulthood. Since expression of GCaMP in Ai93 depends on tTA activity, this manipulation should have prevented GCaMP expression during development, even though doxy-treated mice expressed the same levels of Cre and shared the same genetic background as controls. These experiments suggest that broad expression of GCaMP during development is a major contributing factor to epileptiform activity.

Taken together, these data suggest that a dominant role may be played by GCaMP6 expression itself, perhaps specifically during development. Cre toxicity, tTA toxicity, and/or genetic background, may contribute, but appear unlikely to be determining factors. Whether the GCaMP expression is deleterious only when expressed in certain cell types or brain regions, whether it is deleterious only at certain restricted epochs in development, and whether different versions of the sensor have different severities of effects all remain unknown.

### Available alternative tools

For researchers wishing to pursue experiments with transgenic GCaMP6-expressing mice, some alternatives are available. Pan-excitatory expression with the GCaMP6F sensor can be accomplished with Ai93 using Slc17a7-Cre as an alternative to Emx1-Cre, or with Slc17a7-Cre;Ai95 (with lower expression levels). For the GCaMP6s sensor, the Camk2a-tTA;tetO-GCaMP6s mice (Wekselblatt et al., 2016) have expression levels comparable to Emx1-C re;Camk2a-tTA;Ai94 and have so far (n=12 overall; n=8 with full-hemisphere windows) not been observed to have epileptiform events in our hands.

### Notes of caution

Breeding strategies must take into account the possibility of germline Cre recombination, or else consistent and careful checks must be made to ensure that it has not occurred. This can generally be avoided by excluding females that are positive for both Cre and the *lox*P-flanked allele from breeding pairs. It can be tested for directly by genotyping or by observation, e.g., in the present study, mice negative for Emx1-Cre but positive for Camk2a-tTA and Ai93 (littermates of those positive for all three) would still express GCaMP when germline Cre recombination has occurred.

Finally, though we have identified several lines with apparently normal electrophysiological activity, these mice nevertheless also express the same calcium-buffering proteins and at least one of Cre and tTA. Since a full, precise definition of “normal” and “abnormal” electrophysiological activity is unavailable it is impossible to determine whether all activity in these mice is fully normal. We therefore suggest that confirming observations with multiple techniques – using both transgenic and wild-type mice – is an important experimental control whenever possible.

## Supplemental Movies Caption

A, The fluorescence signal imaged across the dorsal surface of the mouse brain (df/f). B, at top, velocity of a rubber wheel under the forepaws of the mouse; below, traces of df/f over time from the four identified pixels (matching color points in the first panel). Red points indicate detected epileptiform events. C, Two videos of the mouse. All supplemental movies play at half real time. Note that df/f scaling differs between movies for clarity of visualization. For further details of methodology, see Methods of widefield imaging at Carandini/Harris Laboratory.

Movie 1: An Emx1-Cre;Camk2a-tTA;Ai93 mouse.

Movie 2: An Emx1-Cre;Camk2a-tTA;Ai93 mouse.

Movie 3: An Slc17a7-Cre;Ai95 mouse.

## Methods

### Mouse lines

We have used the following abbreviations for transgenic mouse lines (expression notes here are not intended as definitive statements of expression patterns, just as general summaries):

- Ai93 (B6;129S6-*Igs7*^*tm93.1(tetO-GCaMP6f)Hze*^/J, Jax #024103)
  ◦ *Expresses GCaMP6F with Cre and tTA conditionality*
- Ai94 (B6.Cg-*Igs7*^*tm94.1(tetO-GCaMP6s)Hze*^/J, Jax 024104)
  ◦ *Expresses GCaMP6S with Cre and tTA conditionality*
- Ai95 (B6;129S-*Gt(ROSA)26Sor*^*tm95.1(CAG-GCaMP6f)Hze*^/J, Jax #024105)
  ◦ *Expresses GCaMP6F with Cre conditionality*
- Ai96 (B6;129S6-*Gt(ROSA)26Sor*^*tm96(CAG-GCaMP6s)Hze*^/J, Jax #024106)
  ◦ *Expresses GCaMP6S with Cre conditionality*
- Ai32 (B6;129S-*Gt(ROSA)26Sor*^*tm32(CAG-COP4*H134R/EYFP)Hze*^/J, Jax #012569)
  ◦ *Expresses ChR2 with Cre conditionality*
- tetO-G6s (B6;DBA-Tg(tetO-GCaMP6s)2Niell/J, Jax #024742)
  ◦ *Expresses GCaMP6S with tTA conditionality*
- Snap25-G6s (B6.Cg-*Snap25*^*tm3.1Hze*^/J, Jax #025111)
  ◦ *Expresses GCaMP6S in excitatory neurons*
- Camk2a-tTA (B6.Cg-Tg(Camk2a-tTA)1Mmay/DboJ, Jax #007004)
  ◦ *Expresses tTA in excitatory neurons and glia*
- Rosa26-ZtTA (STOCK *Gt(ROSA)26Sor*^*tm5(ACTB-tTA)Luo*^/J, Jax #012266)
  ◦ *Expresses tTA pan-neuronally*
- Emx1-Cre (B6.129S2-*Emx1*^*tm1(cre)Krj*^/J, Jax #005628)
  ◦ *Expresses Cre in excitatory neurons*
- Emx1-Cre-Kess (B6;CBA-Tg(Emx1-cre/ERT2)1Kess/SshiJ, Jax #027784)
  ◦ *Expresses Cre in excitatory neurons*
- Rbp4-Cre_KL100 (here “Rbp4-Cre” - STOCK Tg(Rbp4-cre)KL100Gsat/Mmucd, MMRRC#031125)
  ◦ *Expresses Cre predominantly in isocortical layer 5 excitatory neurons*
- Rorb-IRES2-Cre (here “Rorb-Cre” - B6;129S-*Rorb*^*tm1.1(cre)Hze*^/J,Jax#023526)
  ◦ *Expresses Cre predominantly in isocortical layer 4 excitatory neurons*
- Slc17a7-IRES2-Cre (here “Slc17a7-Cre” - B6;129S-*Slc17a7*^*tm1.1(cre)Hze*^/J,Jax#023527)
  ◦ *Expresses Cre in excitatory neurons, following Vglut1 expression*
- Scnn1a-Tg3-Cre (here “Scnn1a-Cre” - B6;C3-Tg(Scnn1a-cre)3Aibs/J, Jax#009613)
  ◦ *Expresses Cre predominantly in isocortical layer 4 excitatory neurons*
- Cux2-CreER (B6(Cg)-*Cux2*^*tm3.1(cre/ERT2)Mull*^/Mmmh, MMRRC# 032779)
  ◦ *Expresses Cre predominantly in isocortical layer 2/3 and 4 excitatory neurons*
- Ntsr1-Cre_GN220 (here “Ntsr1-Cre” - B6.FVB(Cg)-Tg(Ntsr1-cre)GN220Gsat/Mmucd, MMRC#030648)
  ◦ *Expresses Cre predominantly in isocortical layer 6 excitatory neurons*
- PV-Cre (B6;129P2-*Pvalb*^*tm1(cre)Arbr*^/J, Jax #008069)
  ◦ *Expresses Cre predominantly in fast-spiking inhibitory interneurons*

### Electrophysiology

Electrophysiological recordings were performed at the Carandini/Harris Laboratory (UCL) and were conducted according to the UK Animals Scientific Procedures Act (1986) and under personal and project licenses released by the Home Office following appropriate ethics review. To prepare for electrophysiological recordings, mice were anesthetized briefly (<1 hr) and a small craniotomy was drilled over the site of interest. The craniotomy was covered with Kwik-Cast elastomer for protection and the mouse was allowed to recover. After several hours’ recovery, or over the subsequent several days, mice were head-fixed in the experimental apparatus. The kwik-cast was removed and saline-based agar was applied over the craniotomy. The agar was covered with silicon oil to prevent drying. Recordings were performed with custom multi-site silicon electrode arrays, inserted through the agar and through the dura. Signals were referenced to an external Ag/AgCl wire placed in the agar near the craniotomy. This external reference was shorted to the amplifier ground. LFP signals were low-pass filtered in hardware with a single-pole filter at 300Hz and recorded to disk at 2.5kHz. Subjects were awake and standing passively on a stationary platform. During the analyzed periods, subjects either viewed a “sparse noise” visual stimulus (8-degree width white squares on a black background, at random times and positions) or no visual stimulus. A single channel from those in cortex was selected manually to be representative for further analysis.

### Widefield imaging

At the Carandini/Harris Laboratory (UCL), widefield imaging was performed through the intact skull with a clear skull cap implanted. The clear skull cap implantation followed the method of Guo et al. (2014) but with some modifications. In brief, the dorsal surface of the skull was cleared of skin and periosteum and prepared with a brief application of green activator (Super-Bond C&B, Sun Medical Co, Ltd, Japan). A 3D-printed light-isolation cone surrounding the frontal and parietal bones was attached to the skull with cyanoacrylate (VetBond; World Precision Instruments, Sarasota, FL) and the gaps between the cone and the skull were filled with L-type radiopaque polymer (Super-Bond C&B). A thin layer of cyanoacrylate was applied to the skull inside the cone and allowed to dry. Thin layers of UV-curing optical glue (Norland Optical Adhesives #81, Norland Products Inc., Cranbury, NJ) were applied inside the cone and cured until the exposed skull was covered. A headplate was attached to the skull over the interparietal bone with Super-Bond polymer, and more polymer was applied around the headplate and cone. Imaging was conducted at 35Hz with ~10ms exposures and 2x2 binning using a PCO Edge 5.5 CMOS camera and a macroscope (Scimedia THT-FLSP) with 1.0x condenser lens (Leica 10450028) and 0.63x objective lens (Leica 10450027). Illumination was by 470nm LED (Cairn OptoLED, P1110/002/000). Illumination light passed through an excitation filter (Semrock FF01-466/40-25), a dichroic (425nm; Chroma T425lpxr), and 3mm-core optical fiber (Cairn P135/015/003), then reflected off another dichroic (495nm; Semrock FF495-Di03-50x70) to the brain. Emitted light passed through the second dichroic and emission filter (Edmunds 525/50-55 (86-963)) to the camera. Data were compressed by computing the singular value decomposition of the 3D image stack, and individual pixel traces were reconstructed for analysis from the top 500 singular values.

At the Häusser Laboratory (UCL) all surgical procedures were carried out under a license from the UK Home Office in accordance with the Animal (Scientific Procedures) Act 1986. Emx1-Cre-Kess;Camk2a-tTA;Ai93 mice and Emx1-Cre-Kess;Camk2a-tTA;Ai94 mice, between 6 and 17 weeks old, were anaesthetised using Isoflurane (0.5-1%). A metal headplate with a 5mm circular imaging well was fixed to the skull overlying somatosensory cortex with dental acryclic (Super-Bond C&B, Sun-Medical). A craniotomy was drilled above S1 (right hemisphere, 2mm posterior and 3.5mm lateral of bregma). A cranial window, composed of a 3mm circular glass coverslip glued to a 2mm square glass with UV-curable optical cement (NOR-61, Norland Optical Adhesive) was press-fit into the craniotomy and sealed using Vetbond before fixing it with dental acrylic. After at least 1 week and up to 7 month later widefield imaging was performed at 15Hz using a 470nm LED (Thorlabs, M470L3) to illuminate the area. All except for the first imaging session of cb71-96) were done using an ORCA-Flash 4.0 V3 (Hamamatsu) camera and a 4x objective (4x Nikon Plan Fluorite Imaging Objective, 0.13 NA, 17.2 mm WD). Excitation light passed through an aspheric condenser lens (Thorlabs, ACL2520U-DG15), a filter (Chroma ET470/40) and was reflected into the lightpath by a 495nm longpass dichroic (Semrock, FF495-Di03-25x36) to reach the brain. Emitted light passed through the same 495nm longpass dichroic as well as a 749nm shortpass dichroic (Semrock, FF749SDi01-25x36x3) and an emission filter (Chroma HQ525/50) before reaching the camera. Spontaneous activity of awake mice running freely on a treadmill was acquired for 2-4 minutes. Only the first imaging session of cb71-96 was done using a Manta G609 camera (Allied Vision) and a 50mm f/2.0 Ci Series fixed focal length lens (Edmund optics). Calcium signal time series were obtained from the average pixel intensity.

At the Margolis Laboratory (Rutgers, The State University of New Jersey), widefield imaging was performed through the intact skull that was rendered transparent using methods similar to those described previously (Lee and Margolis, 2016). All procedures were carried out with the approval of the Rutgers University Institutional Animal Care and Use Committee. Mice were anesthetized with isoflurane (3% induction and 1.5% maintenance) in 100% oxygen, placed on a thermostatically controlled heating blanket (FHC) at 36? and mounted in a sterotaxic frame (Stoelting). The scalp was sterilized with betadine scrub and infiltrated with bupivacaine (0.25%) prior to incision. The scalp was reflected and the underlying skull was lightly scraped to detach muscle and periosteum and irrigated with sterile 0.9% saline. The skull was made transparent by applying a light-curable bonding agent (iBond Total Etch, Heraeus Kulzer International) followed by a transparent dental composite (Tetric Evoflow, Ivoclar Vivadent). A custom aluminum headpost was cemented to the right side of the skull and the transparent portion of the skull was surrounded by a raised border constructed using another dental composite (Charisma, Heraeus Kulzer International). Mice were given carprofen (5 mg/kg) postoperatively. Following a recovery period of at least one week, mice were acclimated to handling and head fixation for an additional week prior to imaging. Mice were housed on a reversed light cycle and all handling and imaging took place during the dark phase of the cycle. For imaging, mice were head fixed and the transparent skull was covered with glycerol and a glass coverslip. Imaging was carried out using a custom macroscope that allowed for simultaneous imaging of nearly the entire left hemisphere and medial portions of the right hemisphere. The cortex was illuminated with 460 nm LED (Aculed VHL) powered by a Prizmatix current controller (BLCC-2). Excitation light passed through a filter (479/40; Semrock FF01-479/40-25) and was reflected by a dichroic mirror (Linos DC-Blue G38 1323 036) through the objective lens (Navitar 25 mm / f0.95 lens, inverted). GCaMP6f fluorescence (535/40; Chroma D535/40m emission filter) was acquired with a MiCam Ultima CMOS camera (Brainvision) fitted with a 50 mm / f0.95 lens (Navitar). The resulting 100 x 100 pixel image corresponded to an imaging area of ~6 x 6 mm. Spontaneous activity was acquired in 20.47 s recordings at 100 frames per second with 20 s between recordings. Sixteen movies were acquired in each session. Changes in GCaMP6f relative fluorescence were calculated by subtracting the baseline, defined as the average intensity per pixel in the first 30 or 49 images, from each frame within a movie and then dividing each frame by the same baseline. Calcium signal time series were obtained from the average intensity in 5 by 5 pixel regions of interest.

At the Allen Institute for Brain Science, widefield imaging was performed either through the intact skull using a modification of the method from Silasi et al. (2016), or through a 5 mm diameter chronically implanted cranial window centered over the left visual cortex (Andermann et al., 2010; Zhuang et al., 2017). For through-skull imaging, the skull was exposed and cleared of periosteum and a #1.5 borosilicate coverslip (Electron Microscopy Sciences, #72204-01) was fixed to the skull surface by a layer of clear cement (C&B Metabond, Sun Medical Co, Ltd, Japan). A 3d printed light-shield was fixed around the coverslip using additional Metabond, and the outward-facing surfaces were coated with an opaque resin (Lang dental Jetliquid, MediMark, Grenoble, France). Mice with chronically implanted windows received a 5 mm diameter craniotomy over the left hemisphere, centered at 2.7 mm lateral and 1.3 mm anterior to lambda. The craniotomy was sealed with a stack of three #1 coverslips, attached to each other using optical adhesive (Norland) and to the skull with Metabond. The window provided optical access to the left visual cortex, the posterior aspect of somatosensory cortex and medial aspect of dorsal retrosplenial cortex. In both cases, a custom-manufactured titanium headpost was fixed posterior to the lightshield/coverslip and dorsal to the cerebellum using Metabond. All surgical procedures were performed under isoflurane anesthesia and were approved by the Allen Institute Animal Care and Use Committee. Image acquisition used a Hamamatsu Flash4.0 v2 sCMOS camera running at half resolution (1024x1024) with a 10ms rolling exposure (100Hz). Images were produced by a tandem-lens macroscope (Scimedia THT-FLSP) with 1.0x tube and objective lenses (Leica 10450028) for through skull imaging, or a 1.0x tube lens paired to a 1.6x objective lens (Leica 10450029) for imaging through the chronically implanted window. Epifluorescence illumination used a 470 nm LED (Thorlabs M470L3) filtered (Semrock FF01-474/27-50) and reflected by a dichroic mirror (Semrock FF495-Di03-50x70) through the objective lens. Fluorescence emission passed through a filter (Semrock FF01-525/45-50) to the camera. Data were spatially downsampled 16x by averaging, and calcium traces were obtained by subtracting and then dividing each pixel by its mean over the whole time series.

### 2-photon imaging

The subject (Emx1-Cre;Camk2a-tTA;Ai94, Figure 4) was implanted, the Carandini/Harris Laboratory (UCL), with a 4mm-diameter glass window over visual, retrosplenial, and somatosensory cortices. Two-photon imaging was performed using a standard resonant B-Scope microscope (Thorlabs) equipped with a Nikon 16x, 0.8NA water immersion objective, and controlled by ScanImage 4.2 (Pologruto et al., 2003). Excitation light was provided at a wavelength of 970-980 nm through a tunable Ti:Sapphire laser (Chameleon Ultra II, Coherent). A custom light-isolation cylinder was placed around the imaging objective. Imaging was conducted at 30Hz with a 500µm-wide field of view in superficial layers (depth ~200µm) of somatosensory cortex (a site determined to show epileptiform events in widefield imaging previously). To extract a trace for detection of epileptiform events, all pixels from each frame were averaged together across the entire field of view (in MATLAB: `squeeze(mean(mean(imstack,1),2));` where `imstack` is a 3D matrix of size nXpixels x nYpixels x nFrames).

### Doxycline treatment

Mice treated with doxycycline were administered doxycycline orally via the drinking water. A solution of 5 % sucrose and 2 mg/ml doxycycline in tap water was given as drinking water from birth up until 7 weeks of age. Treated and untreated animals were imaged as described above (Widefield imaging in the Häusser laboratory).

### Testing for Cre-mediated Lox-STOP-Lox recombination in the germline

Mouse DNA was purified from 25 mg tail snips using Macherey-Nagel NucleoSpin Tissue 96 protocol adapted for the QIAgen QIAcube HT platform, yielding DNA concentrations between 15-60 ng/µL as measured by SpectraMax UV Spectrophotometer. Genotyping PCR reaction was performed using primers MG-2333 (5’gctcgtttagtgaaccgtca 3’), MG-1099 (5’ tagccatggtgctgaggggatct 3’), MG-2054 (5’ acagatcccgacccatttgctgt 3’). Amplification was performed in a 50 µL reaction volume and consisting of 9.5 µL nuclease free water, 25 µL Taq master mix, 4.5 µL each primer at 10 µM working concentration, and 2 µL template, and with the following PCR program: 3 min at 95 °C for initial denaturation, followed by 45 cycles of 30 s at 95 °C, 30 s at 60 °C, 90 s at 72 °C, and completed with 10 min at 72 °C. PCR products were visualized by capillary electrophoresis using the Advanced Analytical Fragment Analyzer dsDNA 910 protocol. PCR product size of 119 bp corresponds to intact LSL cassette, PCR product size of 183 bp corresponds to LSL deletion. We observed low-level amplification of 183 bp product in some samples likely due to Cre-mediated recombination in tail tissue. All germline LSL deletion events were observed as complete absence of the 119 bp band.

## Data analysis

Traces were analyzed by finding peaks and computing two parameters from them: prominence and full-width at half-prominence. Prominence is defined as the height of the peak relative to the greater of the two minima between the peak and its surrounding two higher peaks (or beginning/end of trace if no higher peak). Peak detection and calculation of these two parameters can be accomplished with the “findpeaks” function in MATLAB:

`[peakValues,peakLocations,widths,prominences] = findpeaks(signal, t);`

## Acknowledgements

We thank Federico Rossi, Efthymia Diamanti, Sam Failor, Adam Packer, Mehmet Fisek, and Spencer Smith for helpful discussions. We thank Charu Reddy for help with animal husbandry. NAS was supported by postdoctoral fellowships from the Human Frontier Sciences Program and the Marie Curie Action of the EU. CB was supported by an EMBO Long-Term Fellowship and the Marie Curie Action of the EU. MH was supported by the Wellcome Trust, BBSRC and ERC. DJM was supported by the New Jersey Commission on Brain Injury Research, the National Science Foundation and the National Institutes of Health.

